# Molecular evolution and diversity of isomerase–reductase operons involved in bacterial metabolism of glycosaminoglycans

**DOI:** 10.1101/2025.01.30.635655

**Authors:** Yu Nishimura, Kenji Okumura, Sayoko Oiki, Kohei Ogura, Wataru Hashimoto

## Abstract

Glycosaminoglycans (GAGs), comprising uronic acids and amino sugars, are widely distributed in human tissues such as the intestine and oral cavity. Various bacteria colonize these tissues by assimilating GAGs. 4-Deoxy-L-*threo*-5-hexosulose uronate (DHU) is produced during the degradation of GAGs. Pectin, abundant in plants, is also degraded into DHU. DHU is metabolized in a stepwise manner by the isomerase KduI or DhuI, followed by the reductase KduD or DhuD. Previous studies have shown that the two genes encoding isomerase and reductase (*kduI-kduD* and *dhuD-dhuI*, respectively) are designated in an operon. Therefore, it was believed that *dhuD*-*dhuI* and *kduI*-*kduD* operons evolved independently. Nevertheless, the discovery of a hybrid *kduI*-*dhuD* operon raised questions regarding the evolution of these operons. This study was conducted to investigate the factors driving the diversity of operons through a pan-genomic phylogenetic analysis across 3550 bacterial strains. Seven types of DHU metabolism-related operons were identified. The phylum Bacteroidota possesses a hybrid-type *kduI*-*dhuD* operon rather than *kduI*-*kduD* or *kduD*-*kduI* operon. The phylum Bacillota, but not Pseudomonadota or Bacteroidota, possesses the *dhuD*-*dhuI* operon; however, *dhuI*-*dhuD* operon was not detected in any bacterial strain. Although DHU is generated from the degradation of oligomerized GAG by an unsaturated glucuronyl hydrolase (UGL), the UGL gene was found in strains positive for *kduD*-*kduI*, *dhuD*-*dhuI*, *kduI*-*dhuD*, and *dhuD-kduI* operons at high ratios, indicating that the acquisition of these operons is advantageous for colonization on human hosts. (230 words)

**IMPORTANCE:** Glycosaminoglycans (GAGs), crucial components of the extracellular matrix, play a vital role in host infection by pathogenic bacteria as well as in the colonization of the host by commensal bacteria.

The isomerases KduI and DhuI are nonhomologous isofunctional enzymes. The *dhuD*-*dhuI* operon is well conserved within certain phyla and appears to have a strong association with GAG metabolism. In contrast, the *kduI* system is more widely distributed across various species. Based on the possession ratios of genes encoding each enzyme that produces 4-deoxy-L-*threo*-5-hexosulose uronate, this study indicated that the substrates targeted by each metabolic system vary depending on the specific operon type.

## Introduction

Glycosaminoglycans (GAGs) are linear polysaccharides composed of uronic acids and amino sugars and play vital roles in the extracellular matrices of animals (1). Major types of GAGs include nonsulfated hyaluronic acid (HA) (composed of glucuronic acid and *N*-acetylglucosamine), chondroitin sulfate/dermatan sulfate (glucuronic acid or iduronic acid and *N*-acetylgalactosamine with sulfation at different positions), heparin/heparan sulfate (repeating units of uronic acid and glucosamine with varying degrees of sulfation), and keratan sulfate (unique among GAGs as it contains galactose instead of uronic acid, along with *N*-acetylglucosamine) (2).

Specific pathogenic bacteria, such as *Streptococcus pneumoniae* and *S. dysgalactiae*, have evolved specialized gene clusters for the breakdown and utilization of GAGs, especially HA (3–5). This process is crucial for bacterial pathogenicity, because it involves the extracellular breakdown of HA by hyaluronate lyase (HysA), followed by the intracellular transport and degradation of unsaturated disaccharides. GAGs are also important for mediating interactions between the host and gut bacteria (6, 7). The unsaturated disaccharides produced by the lyase reaction are transported into the cytoplasm via the phosphotransferase system (PTS) or ATP-binding cassette (ABC) transporters (4, 8, 9) (Figure 1). In *Streptococcus* species, the disaccharides transported by PTS are degraded by glycoside hydrolase (GH) 88 family of unsaturated glucuronyl hydrolases (UGLs) (8, 10). These UGLs break down the unsaturated disaccharides into unsaturated glucuronic acid and amino sugar. The unsaturated glucuronic acid undergoes a nonenzymatic ring-opening, producing 4-deoxy-L-*threo*-5-hexosulose uronate (DHU). Pectin, a heteropolysaccharide abundant in the plant cell wall, is primarily composed of α-1,4-D-galacturonic acid units; however, it also contains various neutral sugars such as rhamnose, arabinose, galactose, and xylose (11). Pectin lyases are produced by various bacteria such as the plant pathogen *Dickeya dadantii*, the human gut bacterium *Bacteroides thetaiotaomicron*, and the human pathogen *Yersinia enterocolitica* (12). In *D. dadantii*, the pectin oligosaccharides produced by pectin lyases are transported into the periplasm by porins, followed by passage into the cytoplasm through the ABC transporter TogMNAB (13, 14). In the cytoplasm, oligogalacturonate lyases (OGLs) catalyze the conversion of saturated and unsaturated digalacturonate into monogalacturonate and DHU (15). Thus, DHU is produced from GAGs and pectin. DHU is also produced from unsaturated rhamnogalacturonan by unsaturated rhamnogalacturonyl hydrolase (URGH), a member of the GH105 family (16).

**Figure 1.**
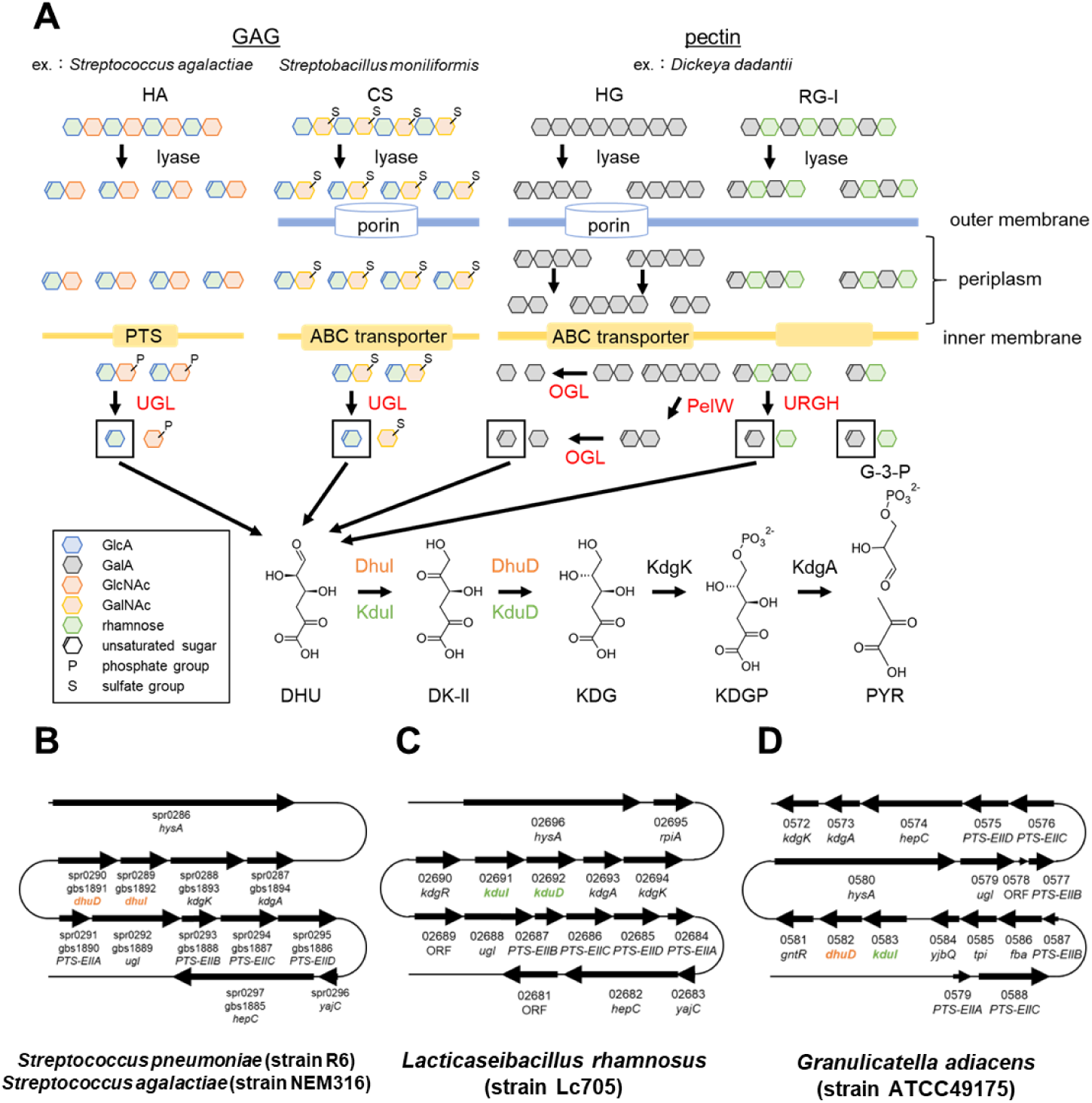
Schematic of metabolism and involved gene clusters. (*A*) Metabolism of glycosaminoglycans (GAGs) and pectins. GAG-metabolizing gene clusters of (*B*) *Streptococcus pneumoniae* (strain R6) (NC_003098.1) and *S. agalactiae* (strain NEM316) (NC_004368.1), (*C*) *Lacticaseibacillus rhamnosus* (strain Lc705) (NC_013199.1), and (*D*) *Granulicatella adiacens* (strain ATCC49175) (GG694015.1). Four or five digits indicate locus tags of each bacterium (spr**** in *S. pneumoniae*, gbs**** in *S. agalactiae*, LC705_***** in *L. rhamnosus,* HMPREF0444_**** in *G. adiacens*).

In *S. agalactiae*, DHU is isomerized by DhuI into 3-deoxy-D-*glycero*-2,5-hexodiulosonate (DK-II), which is further reduced by the NADH-dependent reductase DhuD to form 2-keto-3-deoxy-D-gluconate (KDG) (17). DhuI and DhuD are designated in an operon in the genome of *S. agalactiae* (Figure 1B). The probiotic *Lacticaseibacillus rhamnosus* (strain Lc705) produces the isomerase KduI and the NADH-dependent reductase KduD, whose genes are also designated in an operon as *kduI*-*kduD* (18) (Figure 1C). The operons *dhuI*-*dhuD* and *kduI*-*kduD* encode enzymes that catalyze similar reactions but lack sequence homology, indicating the presence of nonhomologous isofunctional enzymes (NHIEs). KduI and DhuI belong to the KduI/IolB isomerase family (interpro ID, IPR021120) and the sugar-phosphate isomerase, RpiB/LacA/LacB, family (IPR003500), respectively. Both KduD and DhuD belong to the short-chain dehydrogenase/reductase family (IPR002347); however, their sequence identity is low. Therefore, *dhuD*-*dhuI* and *kduI*-*kduD* operons had been believed to evolve independently. Nevertheless, a hybrid type *kduI-dhuD* operon has recently been detected in *Granulicatella adiacens* (previously known as *S. adjacens*) isolated from the oral cavity (Figure 1D) (19). This discovery of the *kduI-dhuD* operon questioned the evolutionary process of operon formation.

Although NHIEs are well-documented, only a few studies have investigated the formation of operons containing NHIEs that code for enzymes catalyzing the same sequential reaction. Furthermore, there has been limited research on the diversity of operon structures across bacterial genomes. Our investigation focuses on operon formation and structure, particularly examining clusters related to the metabolism of GAGs. Hence, the aim of this study was to classify operon structures across bacterial genomes, highlighting the metabolism of GAGs and the intriguing formation of operons involving NHIEs. By understanding these complex genetic arrangements, we aim to clarify the evolutionary pressures shaping bacterial metabolism and operon architecture.

## Results and Discussion

### Variations in isomerase and reductase among whole bacterial strains

There are eight possible combinations (Types A–H) of isomerase (KduI or DhuI) and reductase (KduD or DhuD) (Figure 2). Using representative sequences from 3550 bacterial strains, the DHU isomerases and DK-II reductases present in each strain were identified by BLAST and hidden Markov model search (20, 21). Of the 3550 strains, 562 (16%) harbored the putative operons. From the eight possible types of operon combinations, seven (except Type C: *dhuI-dhuD*) were identified in the bacterial genomes (Figure 3). Among the seven types, strains harboring the *kduI-kduD* operon (Type A) were the most common (280 strains), followed by those harboring the *kduI-dhuD* operon (Type E) (134 strains) (Figure 3B). The *kduD-kduI* operon (Type B), in which the order of the two genes is reversed from that of Type A operon, was widely distributed independent of the phylum as well as Type A operon. The *dhuD-dhuI* operon (Type D) was detected only in the phyla Bacillota (30 strains) and Actinomycetota (6 strains). Moreover, 90% of the strains possessing the hybrid *kduI-dhuD* operon (Type E) belonged to either Bacteroidota (63%) or Bacillota (27%). Type F operon (*dhuD-kduI*) was detected specifically in the phyla Mycoplasmatota and Spirochaetota. Strains harboring the hybrid *dhuI*-*kduD* operon (Type G) were limited to the families Sphingomonadaceae (9 strains) and Erythrobacteraceae (4 strains) in the phylum Pseudomonadota. The *kduD*-*dhuI* operon (Type H) was detected in 41 strains, of which 40 belonged to the phylum Pseudomonadota. Approximately 18% of strains (243 of 1378 strains) belonging to the phylum Pseudomonadota harbored any of Type A–H operons, primarily Types A (13.9%) and H (2.9%). Type H operon was detected only in the phylum Pseudomonadota, except *Faecalibacterium duncaniae* (strain JCM 31915) (phylum Bacillota).

**Figure 2.**
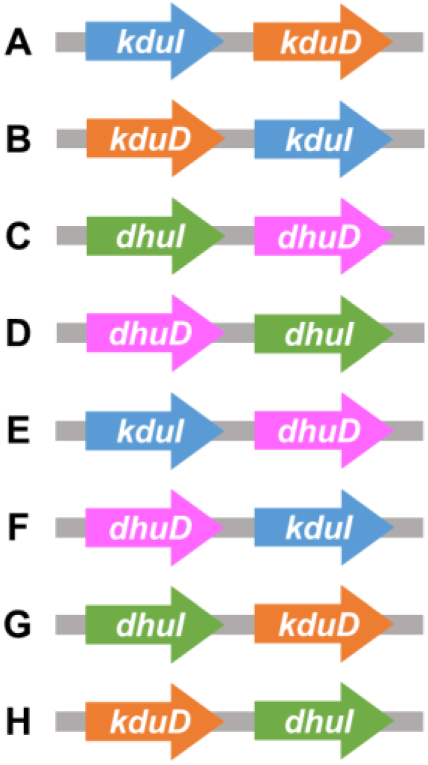
Possible types of isomerase–reductase operons.

**Figure 3.**
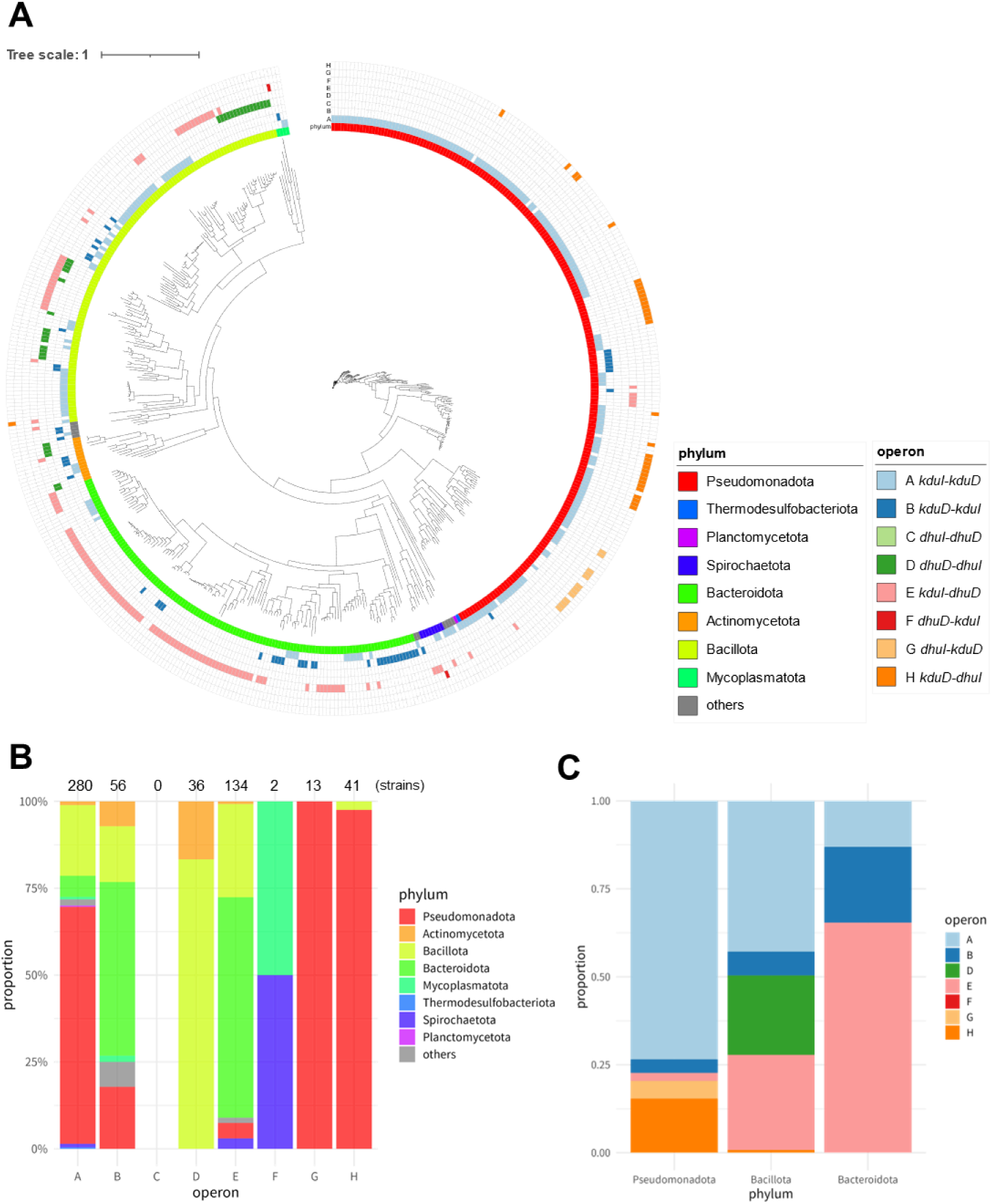
Pan-genome analysis of the operon types. (*A*) A phylogenetic tree based on core genomes and operon types (Types A–H). The figure was drawn using iTOL (version 6) (35). (*B*) Proportions of phyla in each type. The numbers above the bar graphs represent the total number of strains harboring each operon. (*C*) Proportions of the operon types in the phyla Pseudomonadota, Bacteroidota, and Bacillota.

To explore whether the isomerase and reductase are designated as an operon, we calculated the distance between the two genes (Figure 4), which revealed a close distance, except for Type H (*kduD-dhuI* operon). A few Type H strains harbored a gene encoding a cupin-domain-containing protein between *kduD* and *dhuI*. Furthermore, in some strains, *kduI* was positioned upstream of *kduD*, resulting in the formation of the *kduI*-*kduD*-*dhuI* operon. These data suggested that Type H operon was formed through different evolutionary processes.

**Figure 4.**
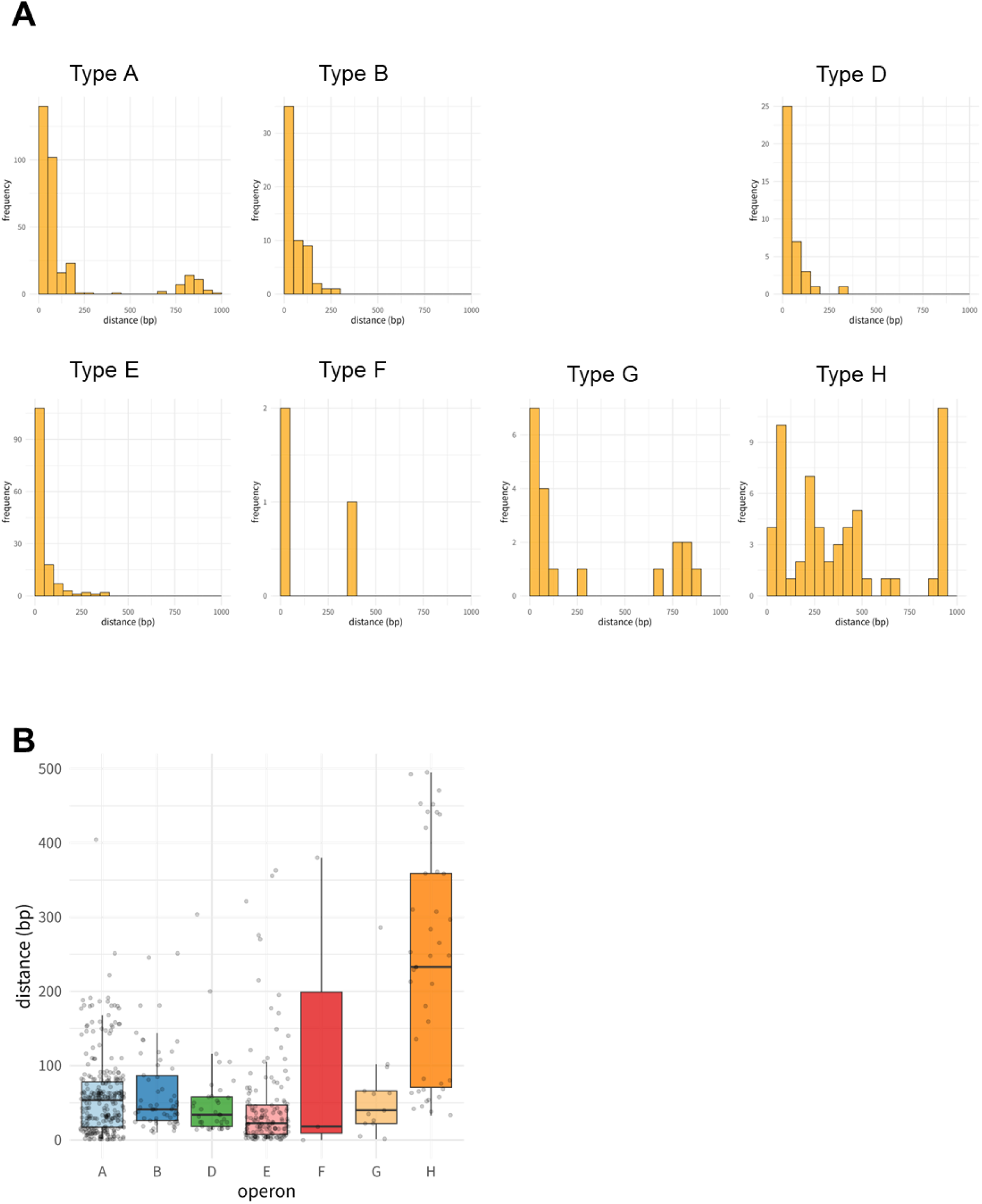
Distance between isomerase and reductase genes. Bars (*A*) and box plots (*B*) were drawn with cut-offs of 1000 and 500 bp, respectively.

### Insights into the phyla Bacteroidota and Bacillota

We further analyzed the diversity of the operons in the phyla Bacteroidota and Bacillota, in which Type E hybrid operon-bearing strains were abundant (Figure 3C and 5). Bacteroidota contained Type A, B, or H operon, whereas one of the strains (*Wenyingzhuangia fucanilytica* strain CZ1127) harbored the tandem operon (*kduD*-*kduI*-*dhuD*). Compared with that in other phyla, the possession ratio of Type A operon was low in Bacteroidota (17 of 309 strains). Bacillota contains the families Streptococcaceae (operon-positive, 15 of 51 strains), Clostridiaceae (operon-positive, 14 of 41 strains), and Lachnospiraceae (operon-positive, 17 of 34 strains), which are commonly resident in the oral cavities and guts of humans and animals. In contrast, no operon-positive strain was detected in the family Staphylococcaceae, which is resident in nasal cavities. Although *Staphylococcus aureus* has been reported to secrete hyaluronate lyases, the lyases are involved in the dispersion of HA-containing biofilms and subsequent distribution of the pathogenic bacterium to host tissues but not in the utilization of HA as nutrient sources (22). Streptococcaceae and Clostridiaceae harbored primarily Type D (*dhuD*-*dhuI*) operons. Five strains in the family Lachnospiraceae (*Marvinbryantia formatexigens* and four *Blautia* species) harbored the tandem *kduI*-*dhuD*-*dhuI* operon.

**Figure 5.**
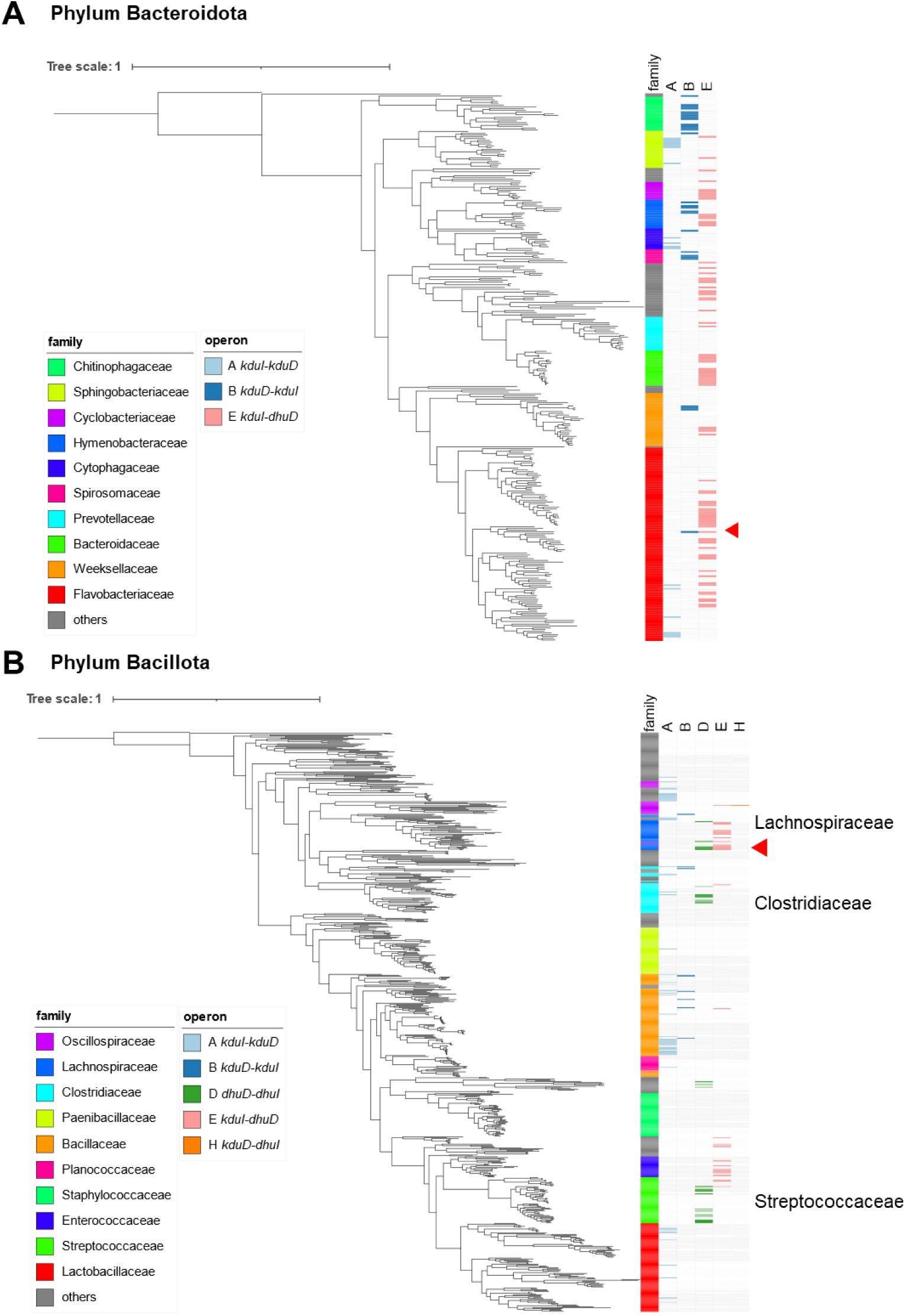
Phylogenetic analysis in the phyla Bacteroidota (*A*) and Bacillota (*B*). The red triangles indicate operons consisting of three genes arranged in tandem. The figure was drawn using iTOL (version 6) (35).

### Correlation with DHU-producing enzymes

Oligomerized GAGs are degraded by UGLs, whereas pectins are degraded by OGLs and/or URGHs. Enzymatic reactions by UGLs, OGLs, and URGHs commonly result in the production of DHU, which are the substrates of the isomerases KduI and DhuI. Therefore, the possession of UGLs, OGLs, and URGHs is suspected to be associated with the environmental adaptation of bacteria. To explore whether the possession of operons correlates with the environmental adaptation, we compared the possession rates of UGL, OGL, and URGH genes among the seven types of operons (Figure 6). Types B-, D-, E-, and F-positive strains harbored UGL genes at high ratios, indicating that the acquisition of these operons is advantageous for colonization on human hosts. Although both OGLs and URGHs target oligomerized pectins, their possession ratios varied in Types D-, E-, and F-positive strains.

**Figure 6.**
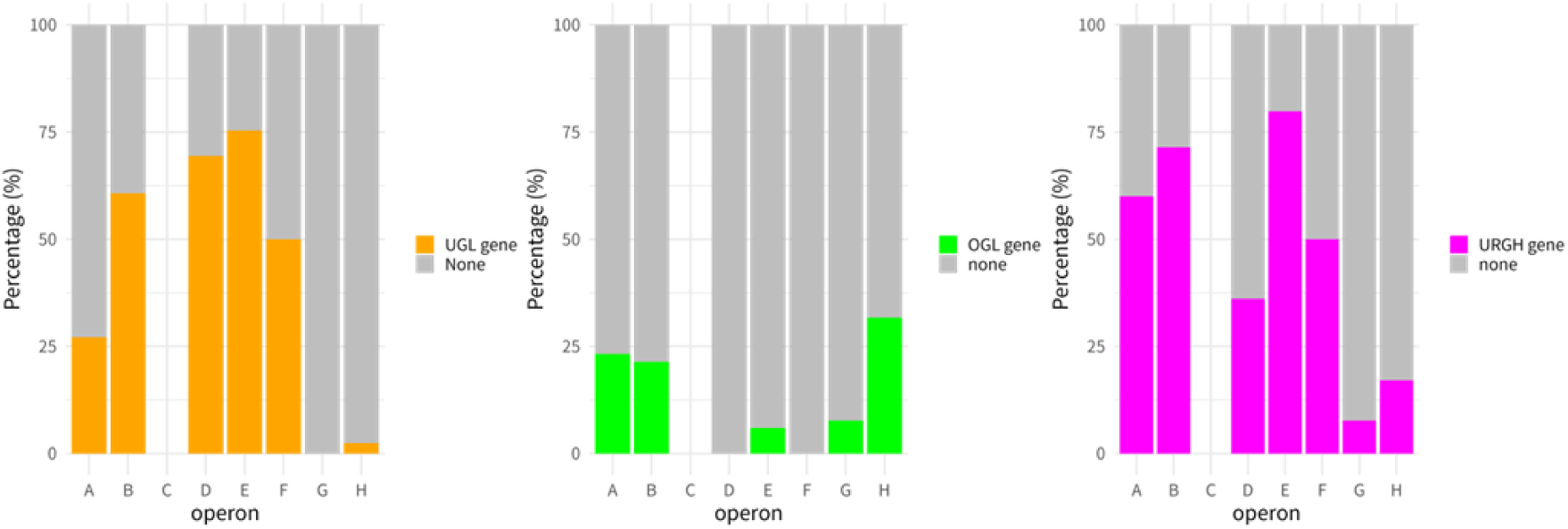
Proportions of UGL, OGL, and URGH gene*-*positive strains among the operon types.

*M. formatexigens* (strain DSM14469), a human gut-commensal anaerobe bacterium, harbors *kduI*-*dhuD*-*dhuI*, which are positive for Types D (*dhuD-dhuI*) and E (*kduI*-*dhuD*). Although the UGL gene was detected in Type D-positive strains, none of them harbored the OGL gene. *M. formatexigens* (strain DSM 14469) harbored a UGL gene but not OGL or URGH gene. However, a previous study demonstrated that this bacterium harbors 103 GHs, including GH28 family hydrolases, which are involved in the digestion of pectin (23), indicating that pectin is degraded into saturated monogalacturonic acid and metabolized independent of KduI or DhuI. Besides *M. formatexigens*, the *kduI*-*dhuD*-*dhuI* operon was detected in four *Blautia* species, including *B*. *producta* (strain DSM 2950), *B. pseudococcoides* (strain YL58), *B. hansenii* (strain DSM 20583), and *B. argi* (strain KCTC 15426). UGL genes were detected in three of the four *Blautia* species (*B*. *producta*, *B. pseudococcoides*, and *B. hansenii*), whereas *B. producta* also harbored URGH, suggesting that the three species can metabolize GAGs. None of the *Blautia* species harbored an OGL gene. It remains unclear whether the genus *Blautia* utilizes pectin as nutrition. Dang *et al.* reported that pectin supplementation to piglets increased the relative abundance of *Blautia* (24). In contrast, Larsen *et al.* demonstrated that pectin supplementation decreased the relative abundance of *Blautia* (25). Because the increase or decrease in the relative abundance does not represent the amount, further analysis is required on the utilization of pectin by *Blautia*.

The family Bacteroidaceae, which contains the genera *Bacteroides* and *Parabacteroides*, harbored only Type E (*kduI*-*dhuD*) operon, which is a hybrid type of *kdu* and *dhu*. *Bacteroides* is abundant in the human gut, which is rich in mucus-derived GAGs and dietary pectins. Therefore, we hypothesized that the possession of Type E operon is advantageous when both GAGs and pectins are abundant. Regarding the phylum Bacillota, Type E operon was detected in the families Lachnospiraceae (16/34 strains), Enterococcaceae (10/23 strains), and Streptococcaceae (3/51 strains). All these Type E-positive species, except the two *Lactococcus* species in the family Streptococcaceae, have been reported as human gut-commensal bacteria (26, 27), supporting our hypothesis. Moreover, UGL genes were detected at the highest ratio in Type E-positive strains (Figure 6). Although the OGL gene was rarely detected in Type E-positive species, they harbored URGH genes at the highest ratio as well as UGL genes. These data indicated that the possession of UGL and URGH genes is advantageous for colonization on the gut as well as the Type E operon.

### Phylogenetic analysis based on isomerase and reductase

We conducted a phylogenetic analysis based on the amino acid sequences of isomerases and reductases (Figure 7). Although a phylogenetic tree of KduI was constructed with five clades (Figure 7A), the clades did not correlate with the types of operons, indicating that variations in KduI sequences occurred before the formation of the operons. In contrast, the variations in Dhu correlated with the types of the operons (Clade 1–Type D, Clade 2–Type G, and Clade 3–Type H) (Figure 7B). This correlation indicated that the variations in DhuI occurred through different selection pressures among the operon-possessing bacteria after the operon formations. DhuI in the phylum Actinomycetota did not form an independent clade but clustered with that of the phylum Bacillota as well as DhuD (Figure S1B). These findings suggest horizontal gene transfer of the *dhuD*-*dhuI* operon from Bacillota to Actinomycetota.

**Figure 7.**
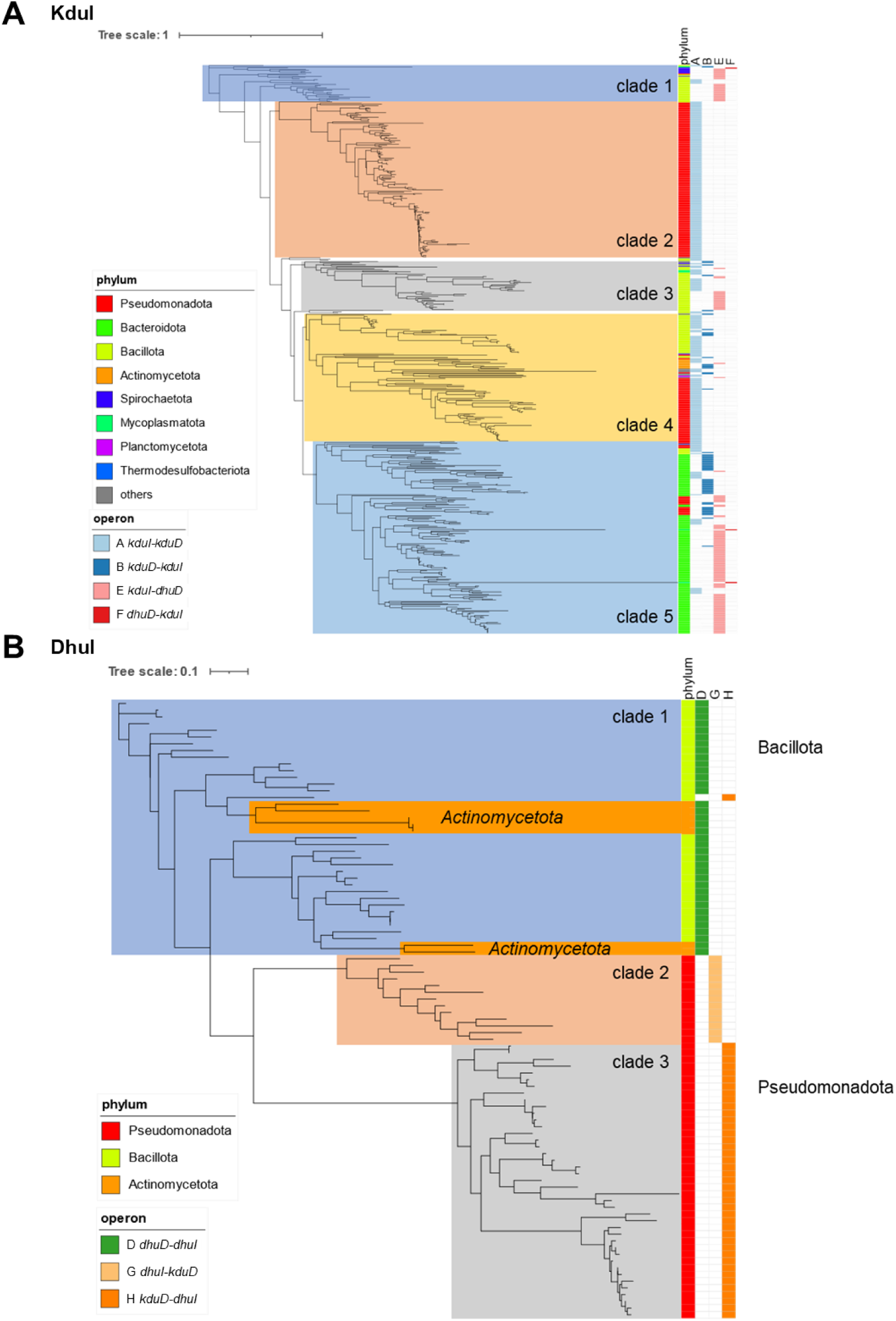
Phylogenetic tree of isomerases (*A*, KduI; *B*, DhuI). The tree was constructed using RaxML (32). The figure was drawn using iTOL (version 6) (35).

In conclusion, this pan-genome analysis hypothesized that Type A operon represents the ancestral form, which was subsequently replaced during the course of evolution. A hybrid operon, Type E (*kduI*-*dhuD*), might play a role in bacterial colonization in the intestinal environment through the metabolism of GAGs and pectins using UGLs and URGHs. Moreover, some species harbored a tandem arrangement of three genes. This study proposes a correlation between Kdu/Dhu genes involved in the assimilation of DHU and evolution of bacterial persistence.

## Materials and Methods

### Data sources

Representative and complete genomes that passed the Taxonomy Check and their annotated protein sequences were obtained from the NCBI RefSeq database (retrieved January 13, 2023) (28). The genomes of 3550 strains were collected, which contained two genomes derived from *Escherichia coli* (O157:H7 str. Sakai and str. K-12 substr. MG1655). Table S1 lists the accession numbers of the genomes. The genomes were classified using TaxonKit (version 0.17.0) (29).

### Phylogenetic analysis

Core genes in the 3550 complete genomes were detected using the UBCG2 pipeline (30). Based on the concatenated sequences of the core genes, a phylogenetic tree was constructed using FastTree (31). Datasets of the phyla Bacteroidota and Bacillota were extracted to construct rooted phylogenetic trees with *Chlorobium limicola* DSM 245 and *Thermotoga maritima* MSB8 for the outgroups of Bacteroidota and Bacillota, respectively. Protein sequences (KduI, KduD, DhuI, and DhuD) were aligned using MAFFT (version 7.525) (32), followed by trimming of poorly aligned regions using the “automated1” method in trimAl (version 1.5.0) (33). The trimmed alignment was used as input for the phylogenetic analysis using RAxML (version 8.2.13) with the PROTGAMMALG model (34). The phylogenetic trees were visualized using iTOL (version 6) (35).

### Homologous gene search based on hidden Markov model (HMM) profile

To generate the HMM profile, 20 nonredundant sequences were selected according to Bundalovic-Torma *et al*. (36) with some modifications. As initial queries, the amino acid sequences of *L. rhamnosus* Lc705 KduI [CAR91530.1], *L. rhamnosus* Lc705 KduD [CAR91531.1], *S. agalactiae* NEM316 DhuI [CAD47551.1], *S. agalactiae* NEM316 DhuD [CAD47550.1], *S. agalactiae* NEM316 UGL [CAD47548.1], *D. dadantii* 3937 OGL [ADM98639.1], and *Bacillus subtilis* subsp. *subtilis* str. 168 YteR [CAB14990.1] were selected (Table S2). Protein BLAST search was conducted using the annotated proteins from the 3550 genomes as references with a threshold of *E* values (<10^−5^) (31). The detected homologous sequences were collected as datasets. Among all the detected homologous sequences, those with sequence identity <97% and with the lowest *E* value were selected as next queries. The sequences of first queries were removed from the datasets. Then, using the next queries and datasets, a protein BLAST search was conducted to detect new queries. By repeating these procedures 20 times, we obtained 21 query sequences for each protein. These selected query sequences were aligned using MAFFT (version 7.525) (32). An HMM profile was generated from the aligned sequences using HMMER (version 3.4) (hmmer.org). Based on the generated profiles, homology searches were conducted using the complete genomes of 3550 strains. Finally, genes whose complete sequences had an *E* value <10^-5^ were considered homologous.

### Identification and classification of GAG metabolism-related gene operons

To determine whether GAG isomerase and reductase are designated in an operon, we calculated the distance between isomerase (*kduI* or *dhuI*) and reductase (*kduD* or *dhuD*) genes detected using the above-described HMMER search. If the distance between the gene pairs was ≤500 bp, the pairs were defined as putative operons. According to the types of genes and order, the putative operons were classified into eight types (Figure 2).

## Acknowledgments

This work was supported in part by KAKENHI (21H02156) from the Japan Society for the Promotion of Science, Japan (W.H.) and by a Research Grant from The Kyoto University Foundation (W.H.). The authors would like to thank Enago (www.enago.jp) for the English language review.

---

**Figure S1.**
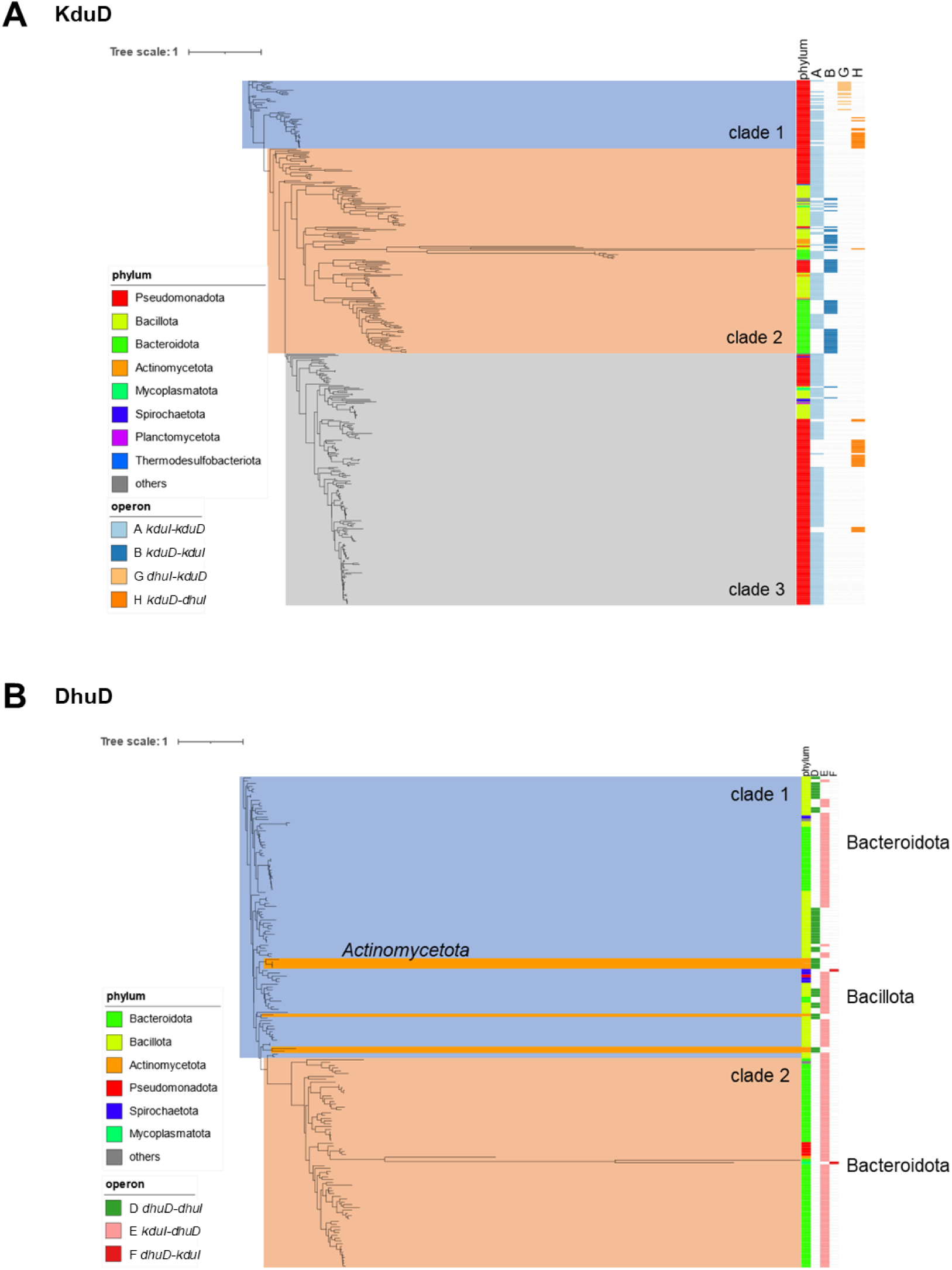
Phylogenetic tree of reductases (*A*, KduD; *B*, DhuD). The tree was constructed using RaxML (34). The figure was drawn using iTOL (version 6) (35).

